# Polyvalent Guide RNAs for CRISPR Antivirals

**DOI:** 10.1101/2021.02.25.430352

**Authors:** Rammyani Bagchi, Rachel Tinker-Kulberg, Tinku Supakar, Sydney Chamberlain, Ayalew Ligaba-Osena, Eric A. Josephs

## Abstract

CRISPR biotechnologies, where CRISPR effectors recognize and degrade specific nucleic acid targets that are complementary to their guide RNA (gRNA) cofactors, have been primarily used as a tool for precision gene editing^1^ but possess an emerging potential for novel antiviral diagnostics, prophylactics, and therapeutics.^2–5^ In gene editing applications, significant efforts are made to limit the natural tolerance of CRISPR effectors for nucleic acids with imperfect complementarity to their gRNAs in order to prevent degradation and mutation at unintended or “off-target” sites; here we exploit those tolerances to engineer gRNAs that are optimized to promote activity at multiple viral target sites, simultaneously, given that multiplexed targeting is a critical tactic for improving viral detection sensitivity,^3^ expanding recognition of clinical strain variants,^6^ and suppressing viral mutagenic escape from CRISPR antivirals.^7^ We demonstrate *in vitro* and in higher plants that single “polyvalent” gRNAs (pgRNAs) in complex with CRISPR effectors Cas9 or Cas13 can effectively degrade pairs of viral targets with significant sequence divergence (up to 40% nucleotide differences) that are prevalent in viral genomes. We find that CRISPR antivirals using pgRNAs can robustly suppress the propagation of plant RNA viruses, *in vivo*, better than those with a “monovalent” gRNA counterpart. These results represent a powerful new approach to gRNA design for antiviral applications that can be readily incorporated into current viral detection and therapeutic strategies, and highlight the need for specific approaches and tools that can address the differential requirements of precision gene editing *vs*. CRISPR antiviral applications in order to mature these promising biotechnologies.

## MAIN TEXT

Class II CRISPR effectors (Figure 1) like Cas9, Cas12, and Cas 13, are nucleases that use a modular segment of their RNA cofactors known as CRISPR RNAs (crRNAs) or guide RNAs (gRNAs) to recognize and trigger the degradation of nucleic acids with a sequence complementary to that segment.^8^ Because of their ability to be easily redirected to nucleic acids with different sequences by simply changing the sequence composition of a short portion of their gRNAs called their ‘spacer,’ CRISPR-based biotechnologies have been rapidly developed over the past several years for a number of different applications, most notably in precision gene editing (Figure 1A).^1^

**Figure 1.**
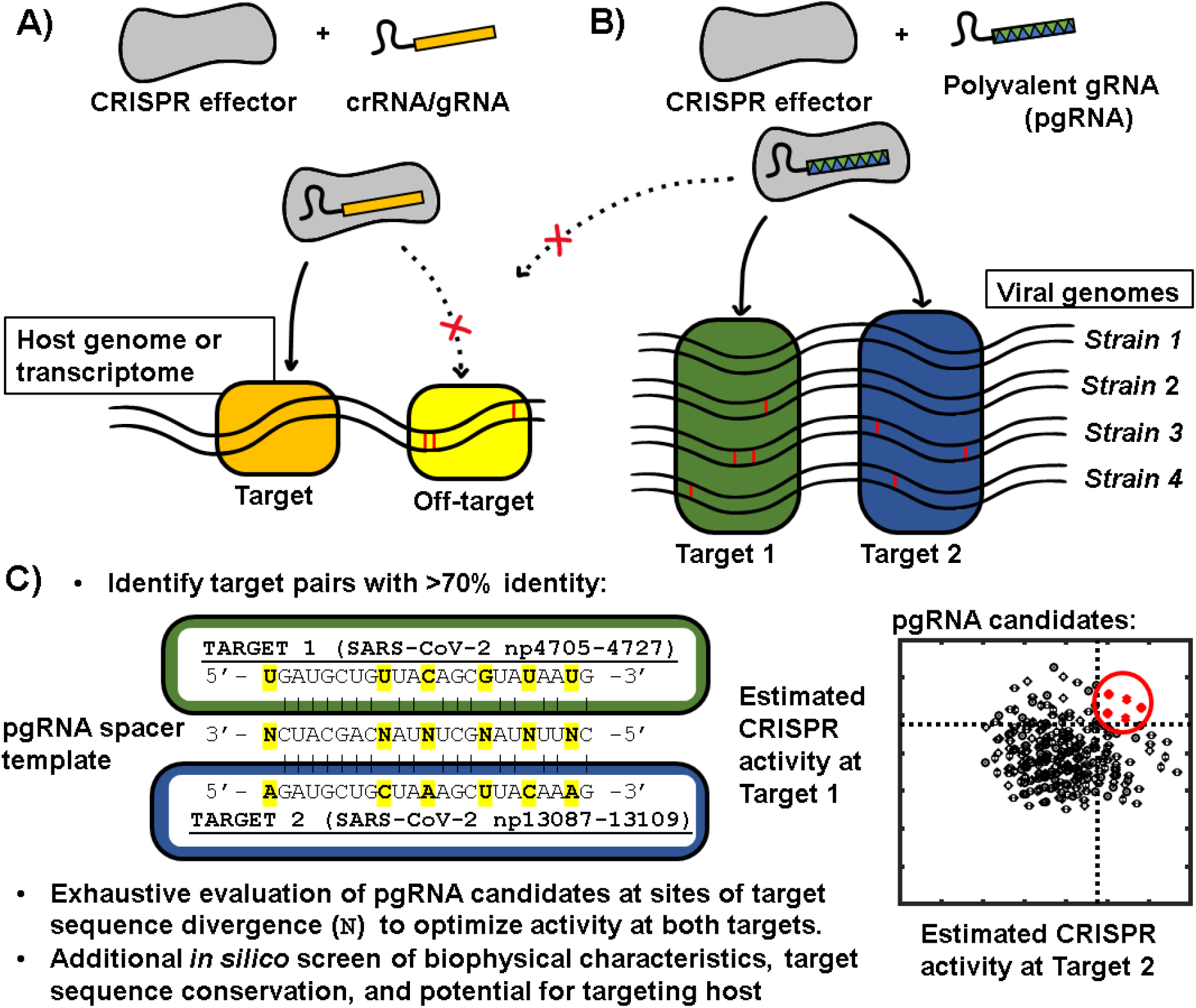
(A) Precision gene editing applications of CRISPR require extreme specificity in mutating only a single target site; however, (B) for antiviral applications of CRISPR, recognition and degradation of multiple viral targets is advantageous for improving viral detection sensitivity, expanding recognition of strain variants, and suppressing mutagenic escape. A single “polyvalent” guide RNA (pgRNA) is optimized to promote activity at multiple viral targets, simultaneously, while avoiding the host genome or transcriptome. (C) Protocol for designing pgRNAs: after target pairs with >70% homology have been identified in the same viral genome, the nucleotides at positions where the sequence between the two targets differ are chosen to minimize potential reductions of activity at the different sites by determining which mismatch- and position-specific mispairings are best-tolerated by the CRISPR effector. See Figure S1 and Supplementary Notes 2 and 3 for details.

Beyond gene editing, another nascent but less-developed application of CRISPR effectors has been as novel antiviral diagnostics, prophylactics, and therapeutics, based on their ability to recognize and degrade viral genomes.^9^ Type II CRISPR effector Cas9 and type V CRISPR effector Cas12, which both recognize and introduce breaks into double-stranded DNA (dsDNA) targets, have been used *in vitro*^7, 10, 11^ and animal models^12^ to degrade dsDNA viruses as well as to excise proviruses from cells with latent retroviral infection.^13^ Because the vast majority of pathogenic viruses are RNA viruses, type VI CRISPR effectors Cas13a (formerly C2c2), Cas13b, and Cas13d, which are RNA-guided RNA endonucleases, also hold significant promise for CRISPR antiviral biotechnologies.^14–17^ Recent demonstrations of the ability of Cas13 variants to reduce viral load by either degrading viral single-stranded RNA (ssRNA) genomes or viral mRNA have been performed in plant,^18–20^ mammalian,^21, 22^ and human^4, 14^ cells, as well as more recently in animal models.^2^ Type V and type VI CRISPR effectors also exhibit nonspecific ssDNAse and ssRNAse activity after recognition of their targets, respectively, and this activity has been harnessed in sensitive strategies to detect viral genetic material in diagnostic devices.^14, 15, 23, 24^ While all of these applications have shown significant promise for the future of CRISPR antivirals, further maturation of these biotechnologies is required before they can reach their full potential.

For example, there are several major challenges to the development of effective CRISPR-based antiviral biotechnologies—challenges which are different from those that arise during the development of effective gene editing biotechnologies—that result from the rapid proliferation and mutation rates of viruses. These include the need to be able to robustly target a specific virus where sequence polymorphisms may exist across strains or clinical variants and, relatedly, the requirement to suppress mutational escape, where the emergence of novel mutations can limit or eliminate the ability of the CRISPR effector to recognize the viral genome. In CRISPR-based antiviral biotechnologies, such issues have been addressed primarily using two approaches: by targeting the CRISPR effector to regions of high sequence conservation in the viral genome,^4, 6, 14^ and by introducing multiple gRNAs to target different segments of the viral genome simultaneously (multiplexing) to make viral escape less likely.^4,14 7^

We propose an alternative strategy for the design of gRNAs for CRISPR antivirals that exploits the widely-recognized tendency of different CRISPR effectors to possess varying levels of tolerance to imperfect complementary between the gRNA spacer and the targets.^25–27^ We hypothesized that, if we could match target sequences in a viral genome to other targets with some shared sequence homology in the same viral genome, a single gRNA spacer sequence could be optimized to maximize CRISPR ribonucleoprotein (RNP) activity at both targets (Figure 1B). This approach is, in effect, the opposite as what is performed during gRNA design for precision gene editing, where significant efforts have gone into limiting this tendency for precision gene editing applications and mutagenic CRISPR activity at multiple or unintended “off-target” sites prevented at all costs (Figure 1A). The development of “polyvalent” gRNAs with one spacer able to target multiple viral target sequences would have multiple advantages for CRISPR antiviral applications: simultaneous targeting of multiple viral sites to increase the probability of generating a disabling mutation and limiting mutagenic escape; reducing the components required for an operative multiplexed targeting to occur; and, for viral detection applications, increasing the effective number of viral “targets” that can be recognized per effector in a given sample to improve detection limits and robustness to clinical variants. However, gRNAs with significant activity at multiple sites would normally be algorithmically rejected by current gRNA design tools that were created primarily for use in gene editing,^26^ so new approaches that are optimized for antiviral applications are necessary (Supplementary note 1).

We developed an algorithm to identify pairs of protospacers (the nucleotide targets of a CRISPR RNP) in the genome of a virus of interest where the target pairs shared at least 70% sequence homology and where levels of CRISPR RNP activity at both targets were computationally predicted to be within the top quartile for all predicted protospacers targeting the viral genome (Figure 1C, S1, and Supplementary note 2). An analysis of 2,372 genomes of RNA viruses in the NCBI Reference Sequence database^28^ revealed that these homeologous pairs of Cas13-targetable sites (23 nt) with >70% identity (>16 out of 23 nt) are prevalent across viral species: RNA viruses with genomes 5,000 nt in length have on average around 30 of such pairs, and those with genomes 10,000 nt in length have on average approximately 120, obeying a power law scaling with genome length (Figure S2, Supplementary Table S1). For human-hosted RNA viruses, we could identify 19,926 of such homeologous target pairs across 89 viruses (Supplementary Table S2).

Candidate pgRNA sequences for each pair were then generated *in silico* by determining what nucleotide positions within the divergent sequences of the two targets would allow for and maximize predicted activity at both sites (Figure 1C), by minimizing the expected “mismatch penalties” or reduction of CRISPR RNP activity at sites with imperfect complementarity to the spacer sequence. Mismatch penalties have been quantitatively determined for several CRISPR effectors: ^25–27,29^ they exhibit a strong dependence on both the type of mismatch (what nucleotides are incorrectly paired) and the position of the mismatch(es) along the target, and have been found to vary not only by type of CRISPR effector but across homologues of the effector derived from different species (Figure S3). For the design of pgRNAs, we computationally maximized the predicted activity of a single gRNA at multiple viral sites by exploiting well-tolerated mismatch- and position-specific mispairings of the CRISPR effectors to minimize potential reductions of activity at the different sites (Supplementary note 3).

As an example of the design workflow, we examined SARS-CoV-2, the coronavirus responsible for the COVID19 respiratory infection, which has a (+) ssRNA genome that is 29,903 nt long (Figure 1C and Figure S4), and toward which Cas13 has recently shown therapeutic potential in epithelial cells and animal models in suppressing viral propagation.^2–4^ We identified 205 pairs of homologous targets (>70% identity) with pgRNA candidates that pass a preliminary biophysical screen (with regards to GC%, secondary structure, and absence of polynucleotide repeats) for a Cas13 RNP. Of those pairs, 144 had pgRNA candidates with predicted activity at both sites ranking in the top quartile of all anti-SARS-CoV-2 gRNAs and no predicted activity *vs*. the human transcriptome (Figure S4A); and of those 144 pairs, 15 pairs (Figure S4B) had perfect sequence conservation at both targeted sites across ∼29,000 sequenced clinical samples (accessed November 23, 2020)^28^ (Supplementary Tables S3 and S4).

To experimentally validate the pgRNA design protocol *in vitro*, we designed pgRNAs for the Cas9 effector from *Streptococcus pyogenes* (SpyCas9), which recognizes and introduces double-strand breaks into dsDNA targets.^13^ These pgRNAs targeted homeologous pairs of synthetic or virally derived protospacers with sequence divergence up to 50%, that is, differing at up to 10 of the 20 bp sites in the protospacers (Supplementary Table 5), and we then measured the cleavage activity of the Cas9 RNPs at those sites *in vitro*. While regular guide RNAs exhibited no cross-reactivity *in vitro* at paired sites with such high sequence divergence (Figures 2A and S5), SpyCas9 RNPs with pgRNAs could consistently cleave both targets even when paired sequences diverged by up to 40% (Figure 2, Figures S6 and S7). In cases where the pgRNA only exhibited activity at one target, those targets could still possess up to 5 mismatches between the pgRNA spacer and the protospacer. Additionally, we found that including a leading 5’-rG on the spacer, a condition thought to result in greater specificity in CRISPR activity for gene editing applications,^30^ consequently reduced pgRNA activity at both sites (Figure S8). These results further highlight the idea that conditions optimized for precision gene editing might not be ideal for maximizing CRISPR activity during antiviral applications (Figure S8). Hence, by optimizing the tolerance for mismatches between the spacer sequence and targeted sites, we show that pgRNAs can be engineered to promote high levels of CRISPR RNP activity at multiple targeted sites simultaneously *in vitro*.

**Figure 2.**
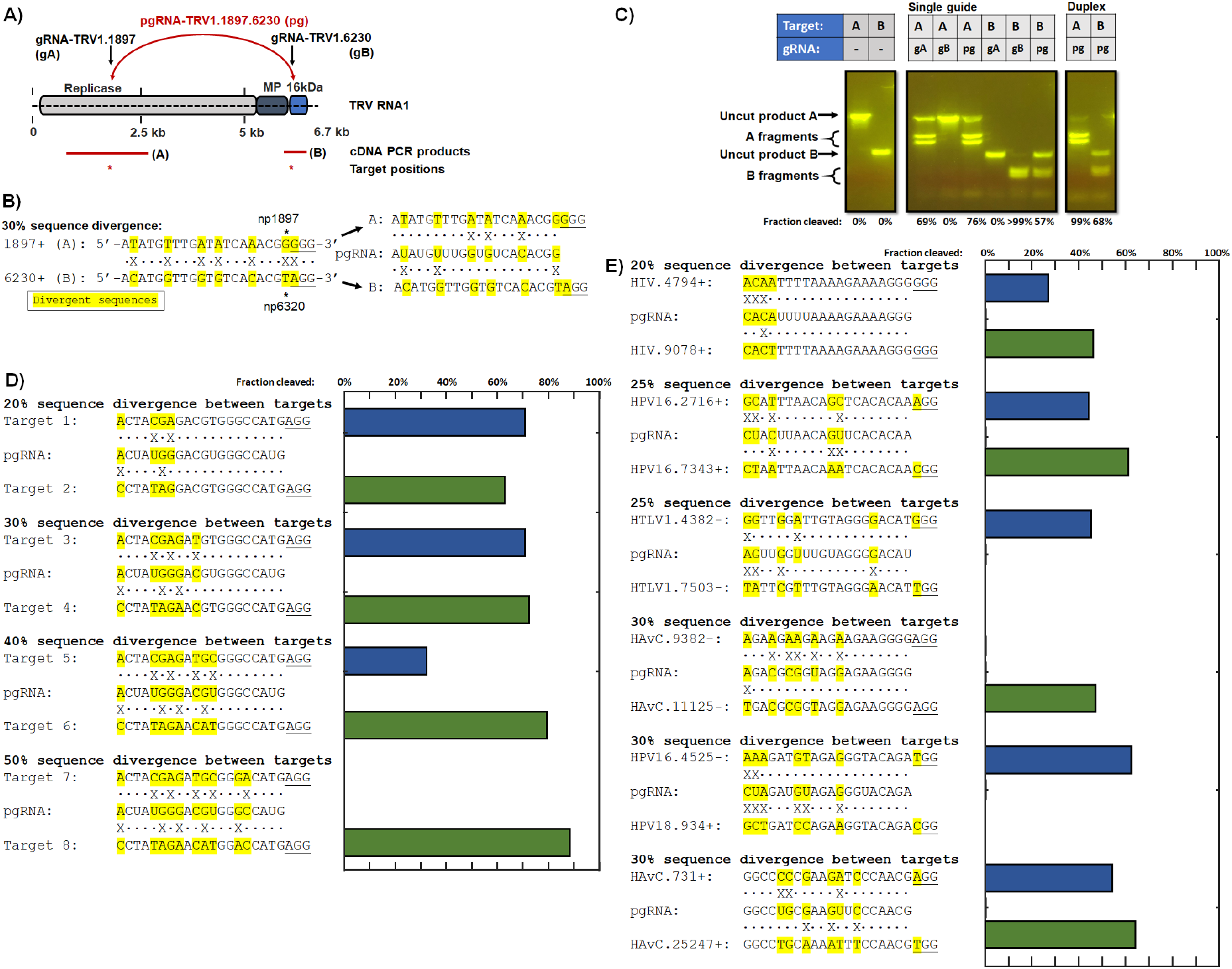
*in vitro* validation of the pgRNA design protocol. (A) A pgRNA (pg) and its two “monovalent” counterpart gRNAs (gA and gB) for SpyCas9 was designed to target two sequences derived from the Tobacco Rattle Virus (TRV) segment 1 (RNA1) at positions 1897 (target A) and 6230 (target B). (B) Targets A and B differ by 6 of the 20 nt (30%) in their protospacer region, and 1 out of 3 within their protospacer adjacent motif (PAM) region (underlined). (C) The monovalent guides exhibit no cross-reactivity at homeologous sites, while the pgRNA exhibits robust cleavage activity at both sites. pgRNA activity is enhanced with a crRNA:tracrRNA duplex compared to a chimeric “single guide” RNA. (D-E) pgRNAs could be generated to exhibit robust cleavage activity at pairs of synthetic (D) and virally derived (E) targets with sequences diverging by up to 40%. HIV: Human Immunodeficiency Virus type 1; HPV16: Human papillomavirus type 16; HPV18: Human papillomavirus type 18; HTLV1: Human T-lymphotropic virus 1; HAvC: Human Adenovirus C.

With pgRNAs, there is a trade-off between potentially decreased activity at a single target and “polyvalent” activity at multiple targets. To test whether pgRNAs could successfully inhibit viral propagation *in vivo*, we designed pgRNAs for RNA-targeting Cas13d from *Ruminococcus flavefaciens* XPD3002 (RfxCas13d) to target tobacco mosaic virus (TMV) replicon (TRBO-GFP) in model tobacco (*Nicotiana benthamiana*) (Supplementary Tables S6 and S7).^20^ RfxCas13d, which has been used in CRISPR-based viral diagnostics^24^ and was recently demonstrated to disrupt influenza and SARS-CoV-2 virulence in human epithelial cells,^4^ is also significantly more tolerant to mismatches between its spacer and target than Cas9 (Figure S3) and does not require specific flanking sequences next to its targets, so RfxCas13d may represent an optimal effector for antiviral applications in that regard.

We introduced a transfer DNA (T-DNA) containing the expression cassette of TRBO-GFP replicon (TRBO-GFP) (Figure 3A) under the control of a constituitive 35S promoter into the leaves of *N. benthamiana via Agrobacterium tumefaciens-*mediated transformation (Figure 3).^18^ The TRBO-GFP replicon lacks a gene for the TMV coat protein, which is required for systemic movement in the plant, but after transcription the (+) ssRNA virus is replication-competent and the virus can spread cell-to-cell within the leaf; a fused reporter gene in TRBO-GFP encoding the green fluorescent protein (GFP) gene allows viral propagation to be tracked visually (Figure 3B and S9). At the same time, we also introduced T-DNAs containing the RfxCas13d gene for transient expression, and T-DNAs to express gRNAs or pgRNAs targeting the viral replicase gene and designed to avoid the *N. bethamiana* transcriptome (transcriptome assembly v5) (Supplementary Tables S7). The plants expressing the “monovalent” counterparts alone (12.6% ± 2.2% (95% confidence) and 22.8% ± 5.5% (95% confidence) GFP mRNA levels relative to plants expressing a gRNA not targeted (NT) to the TRBO-GFP genome (gRNA-NT)) performed less well in viral reduction than the plants expressing the pgRNA, who were able to robustly supress viral spread and viral gene expression (3.4% ± 0.4% (95% confidence) GFP mRNA levels relative to plants expressing gRNA-NT) (Figures 3B-C, Figure S9, and Supplementary Table S8). We hypothesize that the enhanced antiviral suppression observed when using pgRNAs compared to its “monovalent” counterparts is related to the similar improvements found in CRISPR antiviral treatments with multiple, multiplexed gRNAs,^2, 4^ and we will note that pgRNAs can themselves be multiplexed (more than one pgRNA) in antiviral applications to further increase the number of targeted sites per treatment. These results further highlight the potential for CRISPR effectors as viral prophylactic and treatments in plants and other organisms, particularly when combined with pgRNAs specifically optimized for antiviral activities.

**Figure 3.**
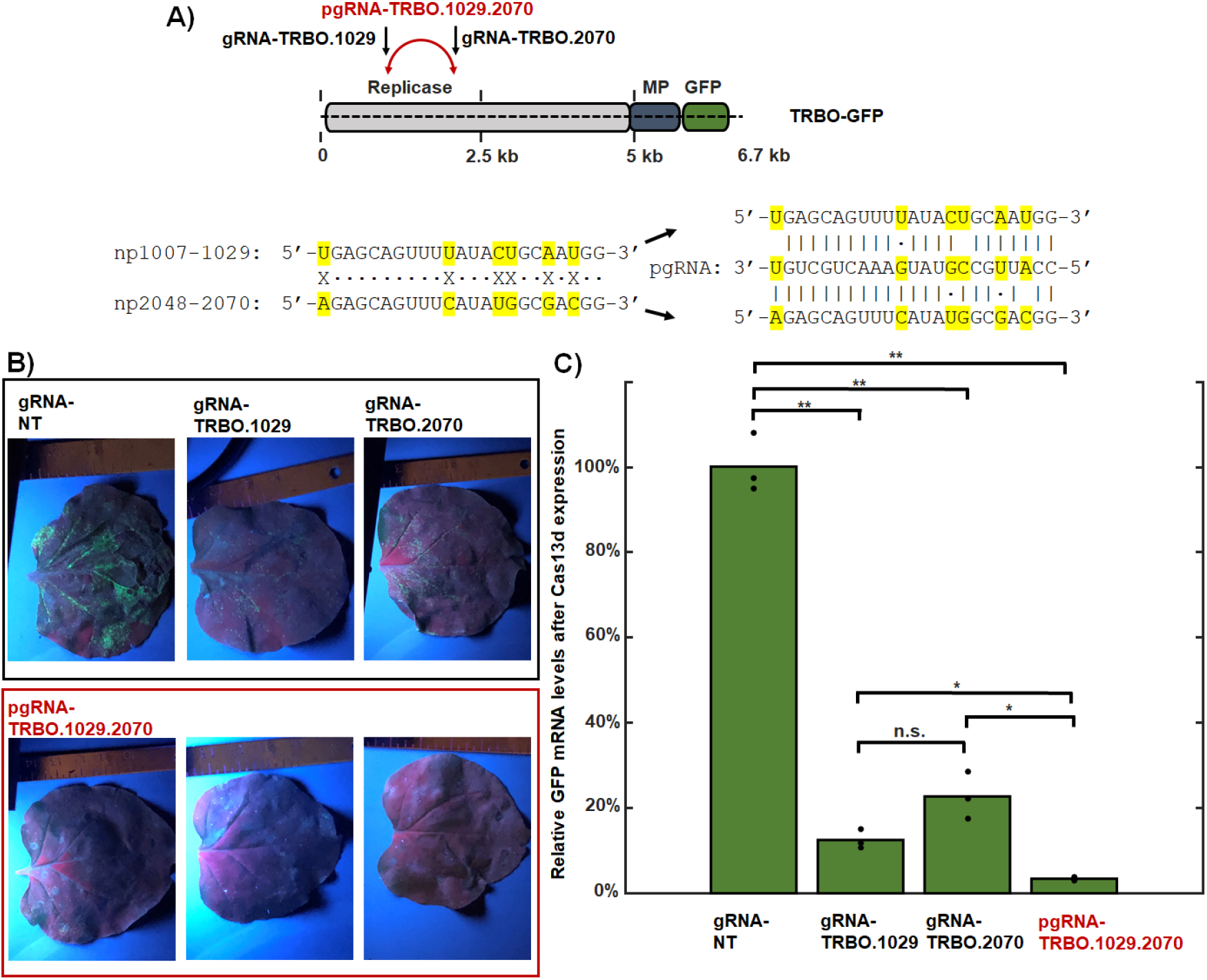
pgRNAs can robustly suppress viral spread in higher organisms (*Nicotiana benthamiana*). (A) A pgRNA for RfxCas13d was designed to target two sequences in a tobacco mosaic virus (TMV) variant replicon (TRBO-GFP) genome. The targets had 30% (6 out of 23) sequence divergence. MP: movement protein. GFP: green fluorescent protein. (B) Leaves were infiltrated with a suspension of *A. tumerfaciens* harbouring plasmids for the transient expression of RfxCas13d; a pgRNA, its two “monovalent” counterpart gRNAs, or a non-targeting (NT) gRNA; and replication-competent TRBO-GFP. Representative images of leaves illuminated under UV light three days after infiltration show the extent of viral spread by GFP expression; see Figure S9 for additional leaf images. Viral spread is suppressed by Cas13 RNPs with gRNAs and strongly by Cas13 RNPs with pgRNAs, but not Cas13 RNPs with a non-targeting (NT) gRNA. (C) Quantitative reverse-transcription PCR (qRT-PCR) of leaf RNA after transient expression demonstrates that pgRNAs are capable of more effectively reducing viral RNA levels than their monovalent counterparts in a higher organism (two-sided T-test; * p < 0.05, ** p < 0.01). N = 4 leaves each for gRNA.

The CRISPR effector proteins used in biotechnological applications were originally found in bacteria and archaea as an antiviral mechanism to degrade foreign DNA and RNA,^8^ and so some tolerance to sequence variation in their targets is likely beneficial for this purpose. In gene editing applications, this tolerance is suppressed to the greatest extent possible using a number of strategies to prevent degradation and mutations at any sequence not exactly matching the gRNA spacer sequence. Here we show that if we identify viral sequences of homeologous target pairs that are abundant in viral genomes then optimize a pgRNA sequence based on the CRISPR effector’s natural position- and sequence-determined tolerance for mismatches, we can achieve robust viral suppression using pgRNAs. In viral detection systems using CRISPR effectors, it has been found that multiplexed use of multiple gRNAs improves viral detection sensitivity,^3^ and we suspect use of pgRNAs for this purpose could similarly improve sensitivity and robustness of viral detection platforms; efforts to adapt the pgRNA design strategy to CRISPR antiviral detection platforms are currently ongoing. While the plant and agricultural applications demonstrated here represent some of the most promising immediate applications of CRISPR antivirals, recent work in human cell and animal models highlight the promise of these approaches as well as the need for gRNA design strategies and CRISPR biotechnologies developed uniquely to satisfy their specific requirements to drive those emerging applications in antiviral diagnostics, prophylactics, and therapeutics.

## CODE AVAILABILITY

Our computational tools for designing pgRNAs for Cas9 and Cas13 are available at https://www.github.com/ejosephslab/pgrna.

## ONLINE METHODS

### Design of polyvalent guide RNAs (pgRNAs)

The pgRNA design algorithm was implemented in MATLAB (Mathworks, Inc.) or Python using code written in-house and made available at: https://github.com/ejosephslab/pgrna. Viral genome sequences used to identify targets are listed in Supplementary Table S9. See Supplementary note 3 for detailed description of the design algorithm.

### Generation of Cas9 target sequences

Cas9 DNA targets in Table 4 were generated by either (1) using plasmids containing targets listed in Table S5 or PCR amplification of targets in plasmids using primers listed in Table S9.

Plasmids were purified using Monarch Plasmid Miniprep Kit following standard protocols (NEB, New England Biolabs, Ipswich, MA). PCR amplification was carried out using 4.5 ng of plasmid DNA, downstream and upstream PCR primers (IDT, Integrated DNA Technologies, Coralville, Iowa, United States) at a final concentration of 0.2uM, and Taq 2x Master Mix (NEB, New England Biolabs, Ipswich, MA, United States) following standard thermocycling protocols. Amplified PCR targets were purified using a Monarch PCR and DNA cleanup kit (NEB) following standard protocols. DNA oligonucleotides were hybridized to form duplex DNA targets by using equal molar concentration of oligos (IDT, Integrated DNA Technologies, Coralville, IA) to a final concentration of 10uM in nuclease-free IDT Duplex buffer. Reactions were heated to 95°C for 2 min and allowed to cool to room temperature prior to the reaction assembly.

### *in vitro* transcription of Cas9 gRNAs

Single guide RNA (sgRNA) was synthesized by using the EnGen sgRNA synthesis Kit (NEB, New England Biolabs, Ipswich, MA, United States) following standard protocols. DNA oligos (IDT) were designed to contain a T7 promoter sequence upstream of the target sequences with an initiating 5’-d(G), as well as overlapping tracrRNA DNA sequence at the 3’ end of the target. The sgRNA was purified using Monarch RNA Cleanup Kit (NEB) and quantitated using standard protocols.

### Duplex gRNA generation

Duplex CRISPR gRNAs (cRNA:tracrRNA) was generated by hybridizing synthetic RNA oligos listed in Table S9 to a universal synthetic tracer RNA oligo (IDT). To hybridize oligos, equal molar concentration of oligos were combined in IDT duplex buffer to a final concentration of 10uM. Reactions were heated to 95°C for 2 min and allowed to cool to room temperature prior to the reaction assembly.

### Cas9 cleavage reactions

Cas9 Nuclease from *S. pyogenes* (NEB) was diluted in 1x NEB Buffer 3.1. prior to the reaction assembly. Cas9 cleavage activity was performed using either PCR-amplified targets, whole plasmid, or hybridized DNA oligos containing desired targets using standard methods. Briefly, Cas9 was preincubated with either a sgRNA or duplex gRNA (crNA:tracRNA) for 5 min at equal molar concentrations in 1x NEB Buffer 3.1 (NEB) in a volume total of 10 ul. Reactions were incubated for 5-10 min at room temperature. Target DNA was then added to the reactions, NEB Buffer 3.1 was added back to a final concentration of 1x, and nuclease-free water was added bringing the final volume to 20 ul. The final reaction contained 100nM Cas9-CRISPR complex and 10nM of target DNA. Similar reactions without the addition of gRNAs to Cas9 were used as a control for uncut DNA. Reactions were incubated at 37°C for 1 hour, followed by the addition of 1 unit of Proteinase K and further incubation at 56°C for 15 min. Reactions were stopped by the addition of one volume of purple Gel Loading dye (NEB).

Fragments were separated and analyzed using a 1.5% Agarose gel in 1xTAE and 1X SYBR Green 1 Nucleic Acid Gel Stain (Thermo Fisher Scientific; Waltham, MA), and fluorescence was photographed and measured (AmershamTM Imager 600; GE Life Sciences, Piscataway, NJ, United States).

### Construction of RfxCas13d for in planta expression

The DNA sequences of the plant codon optimized Cas13d-EGFP with the Cas13d from *Ruminococcus flavefaciens* (RfxCas13d) flanked by two nuclear localization signal (NLS) was amplified from plasmid pXR001 (Addgene #109049) using Q5 high fidelity of DNA polymerase (NEB). Similarly, overlap extension PCR was performed to amplify plant expression vector pB_35S/mEGFP (Addgene #135320) with ends that matched the ends of the Cas13 product so RfxCas13d expression would be under the control of 35S Cauliflower mosaic virus promoter. The PCR products were treated with DpnI (NEB), assembled together in a HiFi DNA assembly reaction (NEB), transformed into NEB10b cells (NEB), and grown overnight on antibiotic selection to create plasmid pB_35S/RfxCas13. Successful clones were identified and confirmed by sequencing followed by transformation into electro-competent *Agrobacterium tumefaciens* strain GV3101 (pMP90).

### Construction of crRNA expression vector

Single stranded oligonucleotides corresponding to “monovalent”, non-targeting (NT), and “polyvalent” gRNAs were purchased from Integrated DNA Technologies (Coralville, IA), phosphorylated, annealed, and ligated into binary vector SPDK3876 (Addgene #149275) that had been digested with restriction enzymes XbaI and XhoI (NEB) to be expressed under the pea early browning virus promoter (pEBV). The binary vector containing the right constructs were identified, sequenced and finally transformed into *Agrobacterium tumefaciens* strain GV3101.

### Agroinfiltration of *Nicotiana benthamiana* (tobacco) leaves

In addition to pB_35S/RfxCas13 and the SPDK3876’s harboring gRNA sequences (TRV RNA2), PLY192 (TRV RNA1) (Addgene #148968) and RNA viruses TRBO-GFP (Addgene # 800083) were individually electroporated into *A. tumefaciens* strain GV3101. Single colonies were grown overnight at 28 degrees in LB media (10 g/L tryptone, 5 g/L yeast extract, 10 g/L NaCl; pH 7). The overnight cultures were then centrifuged and re-suspended in infiltration media (10mM MOPS buffer pH 5.7, 10mM MgCl2, and 200 μM acetosyringone) and incubated to 3-4 hours at 28 degrees. The above cultures were mixed to a final OD600 of 0.5 for CasRX-NLS-GFP-pB35, 0.1 for PLY192 (TRV RNA1), 0.1 for RNA2-crRNAs and 0.005 for TRBO-GFP and injected into healthy leaves of five to six-week-old *N. benthamiana* plants grown under long-day conditions (16 h light, 8 h dark at 24 °C). A total of four leaves for each gRNA were infiltrated. Three days post-transfection, leaves were cut out and photographed under a handheld UV light in the dark, and stored at −80°C before subsequent analysis.

### Quantitative RT-PCR

Total RNA was extracted from infiltrated leaves using RNeasy Plant Mini Kit (Qiagen) and the yield was quantified using nanaodrop. A total of 1ug RNA from control (NT gRNAs) and experimental samples were used for DNase I treatment (Ambion, AM2222) followed by reverse transcription using a poly-dT primer and the Superscript III First Strand cDNA Synthesis System for RT–PCR (Invitrogen). Quantitative PCR was performed on Quant studio 3 Real-Time PCR System from Applied Biosystem using iTaq PowerUP™ SYBR Green pre-formulated 2x master mix (Applied Biosystems). Relative expression levels based on fold changes were calculated using the ddCT method. Cycle 3 GFP mRNA expression levels from the TRBO-GFP replicon were normalized against transcripts of the tobacco PP2A. The samples were performed in three biological replicates.

### Prevalence of pgRNA target pairs in viral genomes

All complete sequences of all RNA viruses with human, mammal, arthropoda, aves, and higher plant hosts found in the NCBI Reference Sequence database were subjected to a brute force direct (nucleotide-by-nucleotide, no gaps) alignment for each of their 23 nt sequence targets to each other, considering only sequence polymorphisms at the same site. We considered only the (+) strand, as even for (-) and dsRNA viruses these sequences would match the vast majority of mRNA sequences. Only targets lacking polynucleotide repeats (4 consecutive rU’s, rC’s, rG’s, or rA’s) were considered viable targets. Targets derived from different segments or cDNAs of the same viral strain were considered together. In total: arthropoda (1074 viral species), aves (111), mammal (496), higher plant / embrophyta (691), and human (89) - hosted viruses were considered (Supplementary Tables S1 and S2).

### Estimation of SARS-CoV-2 target sequence conservation

All complete SARS-CoV-2 genomic sequences available from the NCBI Virus database were downloaded on November 23, 2020 (29,123 sequences). For each of the 205 target pairs possessing biophysically feasible pgRNA candidates, we aligned (no gaps) each target sequence to each genome to determine the closest matching sequence. Alignments containing ambiguous nucleotide calls were not included. Sequence variants were grouped together, with a minimum prevalence of 0.1%, with the fraction of hits by the most prevalent group being considered the sequence conservation reported.

## SUPPLEMENTARY DATA

Supplementary Data are available online.

## CONTRIBUTIONS

EAJ conceived the project. RB, RTK, AL-O, and EAJ designed the experiments. RB, RTK, and AL-O performed the experiments. EAJ and TS wrote the computational code for pgRNA design. EAJ and SC performed the analysis of pgRNA target pair prevalence in viral genomes and of SARS-CoV-2 pgRNAs. RB, RTK, AL-O, and EAJ analysed the data. EAJ wrote the manuscript with contributions from all the authors.

