## Supplemental Data for "Polyvalent Guide RNAs for CRISPR Antivirals"

### SUPPLEMENTARY DATA

**Supplementary Table S1: Statistics of Homeologous Cas13 Target Pair Prevalence (>16/23 or 70% sequence identity) in RNA viral genomes**

**Supplementary Table S2: Homeologous Cas13 Target Pairs (>16/23 or 70% sequence identity) of Human-hosted RNA viruses**

**Supplementary Table S3: RfxCas13d pgRNA Target Pairs and Sequence Conservation for SARS-CoV-2**

**Supplementary Table S4: RfxCas13d pgRNAs for SARS-CoV-2 and Associated Properties of the pgRNAs**

**Supplementary Table S5: Cas9 gRNAs, pgRNAs, and targets for *in vitro* validation of the pgRNA design protocol**

**Supplementary Table S6: RfxCas13d pgRNAs for TRBO-GFP and Associated Properties of the pgRNAs**

**Supplementary Table S7: RfxCas13d gRNAs and pgRNAs used in *N. bethamiana***

**Supplementary Table S8: RT-qPCR of GFP mRNA in *N. bethamiana* after transient expression of TRBO-GFP; RfxCas13d; and gRNAs, pgRNAs, or non-targeting (NT) gRNAs**

**Supplementary Table S9: Oligos, DNA, and Genomic sequences used in this study**

**Supplementary note 1:** Similarities and differences in the design criteria for gRNAs used for precision gene editing and those used for CRISPR antivirals

Despite significant differences in the goals and desired outcomes between CRISPR precision gene editing and CRISPR antivirals (Figure 1), there are some primary objectives in the design of targeting sequence of the gRNA spacer sequences shared by both applications. In particular, CRISPR activity at the desired target is maximized by identifying spacer sequences with no or weak internal secondary structures, moderate GC content (GC%, between ~30%-70%), avoidance of polynucleotide repeats that may inhibit gRNA expression, and avoiding chromatin[1] or occluded targets. Recent bioinformatic analyses have revealed additional sequence contexts and features that may be used to predict spacer sequences with maximized on-target CRISPR activity [2-4].

In the case of precision gene editing (Figure 1A), avoidance of CRISPR activity at any unintended or 'off-target' site is of paramount importance to prevent unwanted genetic mutations [5]. When some flexibility exists in the choice of a specific target (the mutational knockout of a gene, for example), this is achieved by designing gRNA spacers targeted to sites with few other similar genomic sequences, or is otherwise performed by increasing the specificity of the CRISPR effectors, that is, limiting the tolerance of CRISPR effector for any mispairs between the spacer and the target [6-8]. Increasing specificity of CRISPR systems for gene editing applications has been the subject of significant efforts, from structure-based engineering[3, 9] and directed evolution[10] of the CRISPR effectors themselves to the destabilization or fine-tuning of spacer-target interactions to limit activity at sequences that are similar but imperfect matches to the desired target.[11, 12]

In contrast, for CRISPR antivirals (Figure 1B), avoidance of activity with human genomic or transcriptome must be balanced against a requirement for tolerance to sequence heterogeneities across viral targets. In antiviral applications, of paramount importance is the prevention of mutagenic escape—the loss of CRISPR antiviral activity as a result of heterogeneity across clinical strains or viral families at the target site, or as a result of non-inactivating mutations that might occur after mutagenic repair of CRISPR degradation at the target [13, 14]. During antiviral applications, these challenges have typically been addressed by simultaneously introducing multiple gRNAs (up to six) to target different regions of the viral genome, limiting the possibilities for mutagenic escape, and targeting regions of high sequence conservation or functional importance where mutations might not be well tolerated [15-17]. Currently, the design of gRNAs for CRISPR antivirals relies on the computational tools used for precision gene editing, which may lead to sub-optimal antiviral outcomes.

**Supplementary note 2:** Design principles for polyvalent gRNAs (pgRNAs)

The design of polyvalent gRNAs or pgRNAs relies on exploiting known tolerances of CRISPR effectors for mismatches between gRNA and the target to maximize activity at multiple viral sites [2, 4, 5, 18-20]. These tolerances exhibit a strong dependence on both the type of mismatch (what nucleotides are incorrectly paired) and the position of the mismatch(es) along the target, and vary not only by type of CRISPR effector but across homologues of the effector derived from different species (Figure S3).

Careful and systematic studies have been performed to better predict and minimize the propensity of "off-target" effects gene editing; for the design of pgRNAs, we can use these same studies to instead attempt to maximize activity of a single gRNA at multiple viral sites. A metric to score the relative propensities of a CRISPR effector at a site that does not perfectly match its target that is both powerful and simple-to-implement uses a Cutting Frequency Density (CFD) [4, 5, 19] matrix to estimate the penalty or relative decrease in CRISPR activity at off-target sites as a result of each difference in sequence between the target and that site. This approach is described in more detail in Supplementary Note 3. The CFD matrix consists of the mismatch- and position-specific penalties that have been derived from massively parallel characterizations of off-target CRISPR activity, and for each expected mispairing between the gRNA and the off-target site, these penalties are multiplied together to obtain a final score or relative expected CRISPR activity at that site. CFD scores in precision gene editing are used to reject gRNAs which may exhibit high activities at multiple sites in a targeted genome.

The design of pgRNAs uses CFD scores to maximize predicted activity at multiple viral sites following this approach (Figure S1): (i) the activity for each potential target in a virus is predicted using an algorithm like sgRNA designer or cas13design for Cas9 or Cas13, respectively, and those sites with guides ranked in the top quartile are identified and matched those with sequence similarity (*e.g.*, >16 out of 23 nt sequence identity); (ii) the positions of sequence differences between the pairs are located; (iii) a "template" pgRNA spacer is generated that is complementary to the shared nucleotide sequences of the targets, and from the template "candidate" pgRNA spacers with different nucleotides at the positions of sequence divergence are created; (iv) the different candidates are then scored according to the CFD at both targets; (v) then, if a candidate receives a passing score (expected relative activity at both sites greater than a threshold level), a further analysis of those candidates is performed. Candidate pgRNAs are evaluated *in silico* for biophysical characteristics, like GC%, secondary structure free energy, and the ability of the 'direct repeat' segment of the gRNA to form (which is essential for CRISPR activity [2, 4]) as preliminary indicators for a high likelihood of strong on-target activities. Further analysis also includes scoring the potential off-target activity in the host genome or transcriptome and determining the minimum relative activity across variants (MRAV) by calculating CFD for the pgRNA candidates at each site across different clinical viral strains (tolerance to sequence heterogeneities). In this way, our gRNA design algorithm focuses explicitly on the major design considerations (multiplexing/preventing escape; tolerance for clinical variation / viral sequence heterogeneity) for CRISPR antivirals applications.

**Supplementary note 3:** Detailed description of pgRNA design algorithm

#### Calculation of mismatch penalties and relative CRISPR activities

Estimates of the relative CRISPR activity at sites not perfectly targeted by the gRNA/pgRNA spacer sequence were generated by calculating the Cutting Frequency Determination (CFD) score.[5, 18] To calculate the CFD score, the penalty (relative reduction in CRISPR activities) that result from each site with a mismatch is first drawn from a CFD matrix, the table of position-specific reductions of activity that occur as a result of mispairing between specific nucleotides in the spacer and target. The CFD matrices for CRISPR effector were generated by the Sanjana lab (RfxCas13d[2]) and Doench lab (SpyCas9[5, 18], using the data from the "dropout" experiments) using massively parallel screens of gRNA libraries for CRISPR activity, and CFD scoring implemented using publicly available data sets from those labs. The CFD score for a given target and gRNA spacer is the product of the CFD penalties for each mismatch; the position-specific penalties (average over all possible mismatched nucleotides) are summarized in Figure S3. This approach is fast to implement and has been successfully used as a reasonable approximation for CRISPR activity at off-target sites by for a number of different CRISPR effector.[18, 19]. In the case of RfxCas13d, penalties were recovered from taking the value of the reported log2(Fold-Change in expression) to the second power, vs. a perfectly complementary targeted mRNA reporter in their massively parallel screen for gRNA activity in the presence of mismatches.[2] A missing value (rA-rC mismatch at position 15) was interpolated from the penalties of the rA-rC mismatches at positions 14 and 16. In the event of multiple sequential mismatches (two-in-a-row, three-in-a-row, etc.), the position-specific penalties for double- and triple- mismatches were used to calculate the CFD scores at those sites. If the off-target sites had <15 nt identity as the intended target (<55% identity), the CRISPR effectors were considered effectively inactive at those sites. In the case of Cas9, changes in the relative activity at sites with non-canonical PAM sequences (apart from the canonical PAM sequence d(NGG)) are also included as part of the CFD score.

#### Design of polyvalent guide RNAs

The protocol for the design of polyvalent guide RNAs is summarized in Figure S1, and implemented using MATLAB R2018a (Natick, MA) with the Bioinformatics Toolbox or Jupyter notebook with the biopython package and the NCBI-BLAST+ suite.[21] To elaborate on each step of the protocol:

*Step 1: Estimate activity at different targets ('protospacers')*. "On-target" activity for every potential target in a viral genome is found using sgRNA Designer for Cas9[22, 23] or cas13design for Cas13d,[2]software. Only those targets with predicted activity in the top quartile are generally considered as potential pgRNA targets.

*Step 2: Identification of Targetable Pairs with high homology.* Every potential target is aligned to every other potential target, and pairs with >70% sequence identity (≥ 14 nt identity for 20 nt Cas9 targets and ≥ 16 nt identity for 23 nt Cas13d targets) are identified.

*Step 3: Optimization of pgRNA activity at pair sequences.* For a given target pair, a pgRNA spacer template was generated complementary to the targets, using the location and sequences of the matching targets. Different 'candidate pgRNA' spacers were generated with all four potential nucleotides (rA, rU, rC, rG) at each of the sites of sequence divergence between the target pairs, *i.e.* $4^n$ candidates for target pairs with n differences between sequence. A mismatch penalty (CFD score) between the candidates and each of the target pairs was calculated using the multiplicative approach (*i.e.*, Figure 1C right). Those with activity at both sites predicted to remain in the top quartile (or other threshold) for both were kept for further evaluation. Candidate pgRNAs with homopolymer repeats (≥4 consecutive 'rU' or ≥5 consecutive 'rG', 'rC', or 'rA') were removed. Those with GC% <30% or >70% were also removed from consideration. For RfxCas13d, the respective 'direct repeat' sequence for each crRNA (5′-ACCCCUACCAACUGGUCGGGGUUUGAAAC-3′) sequence was appended 5'- to their pgRNA candidate spacers and the pgRNA secondary structures evaluated using the RNAfold function from MATLAB's Bioinformatic Toolbox or Vienna RNA.[24] If the secondary structure of the direct repeat was perturbed by presence of the candidate spacer from its canonical structure, it was removed from consideration, as were those with secondary structure free energy in the spacer region lower than -5 kcal/mol.

*Step 4: Estimate activity at potential host off-targets.* Candidate pgRNA spacers are aligned to the host genome for Cas9 (*i.e.,* Genome Reference Consortium Human Build 38, GRCh38 human reference genome): in this manuscript *N. bethamiana* transcriptome (transcriptome assembly v5) using a local nucleotide BLAST optimized for short sequences <30 nt (blastn-short). The region surrounding each hits to the human genome or transcriptome, to a total of 23 nt (the 23 nt protospacer for Cas13d and 20 nt protospacer + 3 nt PAM for Cas9), were evaluated for a mismatch penalty score with its respective pgRNA candidates and, for Cas9, the presence of PAM. Those with no predicted interaction with the host genome or transcriptome are considered the leading candidates.

*Step 5: Selection of pgRNA based on additional functional criteria.* At this stage, the pgRNA candidates have been screened for high activity at multiple viral targets, no predicted activity at host "off-target" sites, and biophysical characteristics that suggest they would retain  high overall CRISPR activity.[2, 5] The candidates can then be further refined by considering pgRNA targets located within specific genes or regions of interest (ROIs) that may be of clinical or functional significance, or conservation of the targets / viral intolerance to mutations, prior to experimental validation.

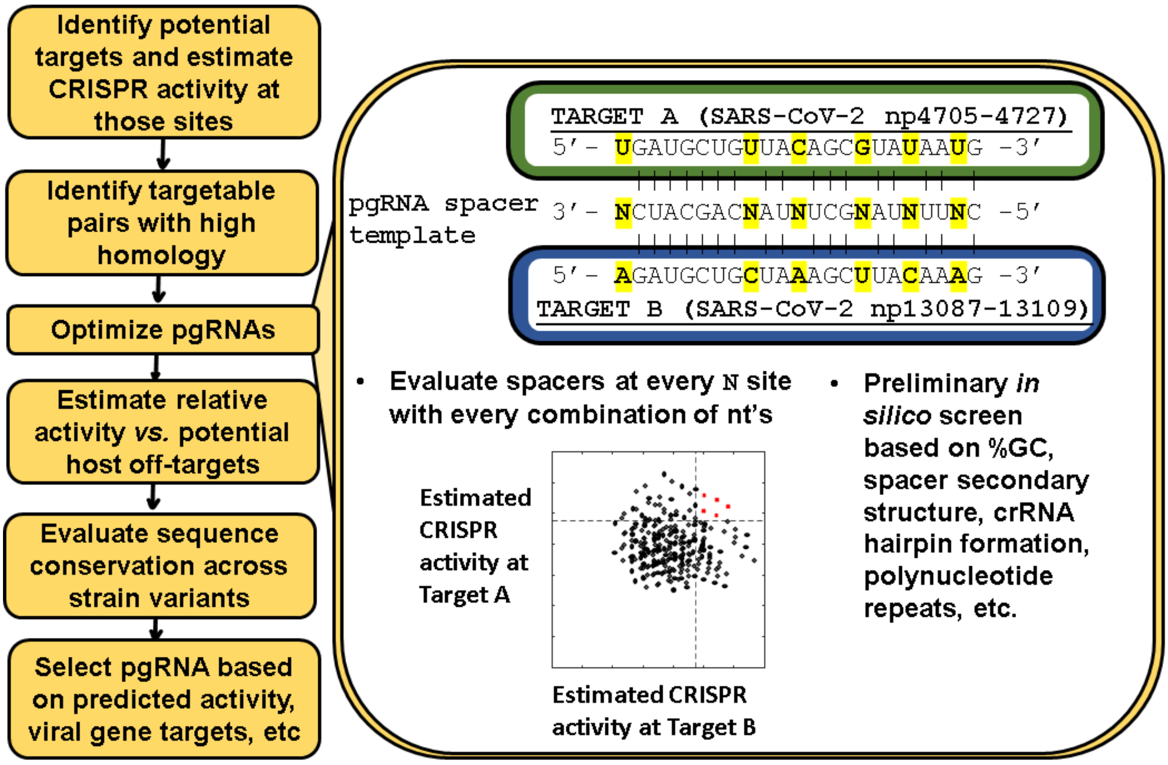


**Figure S1.** Workflow for the computational design of polyvalent guide RNAs (pgRNAs). Briefly, after all possible targets in the viral genome are evaluated for CRISPR predicted activities, using tools such as sgRNA Designer for Cas9[22, 23] or cas13design for Cas13d,[2] pairs of targetable sequences in the top quartile of predicted on-target activity with large fractions of identical sequence (*e.g.*, ≥70%) are identified. A pgRNA spacer template is generated (right). For pairs with n sites where the sequence differs, 4n candidate pgRNA spacers are generated with every possible combination of nucleotides at those n sites, which are then evaluated for sufficient predicated relative activities at both target pairs using a Cutting Frequency Determination (CFD) score. They are then screened in silico for acceptable biophysical properties known to affect CRISPR activity (secondary structure, GC%, etc.). Those pgRNA candidates with acceptably high relative activity across all clinical strain variants and acceptably low predicted activity at potential off-target sites with the human genome/transcriptome can then be further screened for additional criteria (targeting specific genes or regions of interest (ROIs), for examples) and evaluated using additional gRNA design tools or validated experimentally. Implementation of this algorithm in Python is available at: https://github.com/ejosephslab/pgrna.

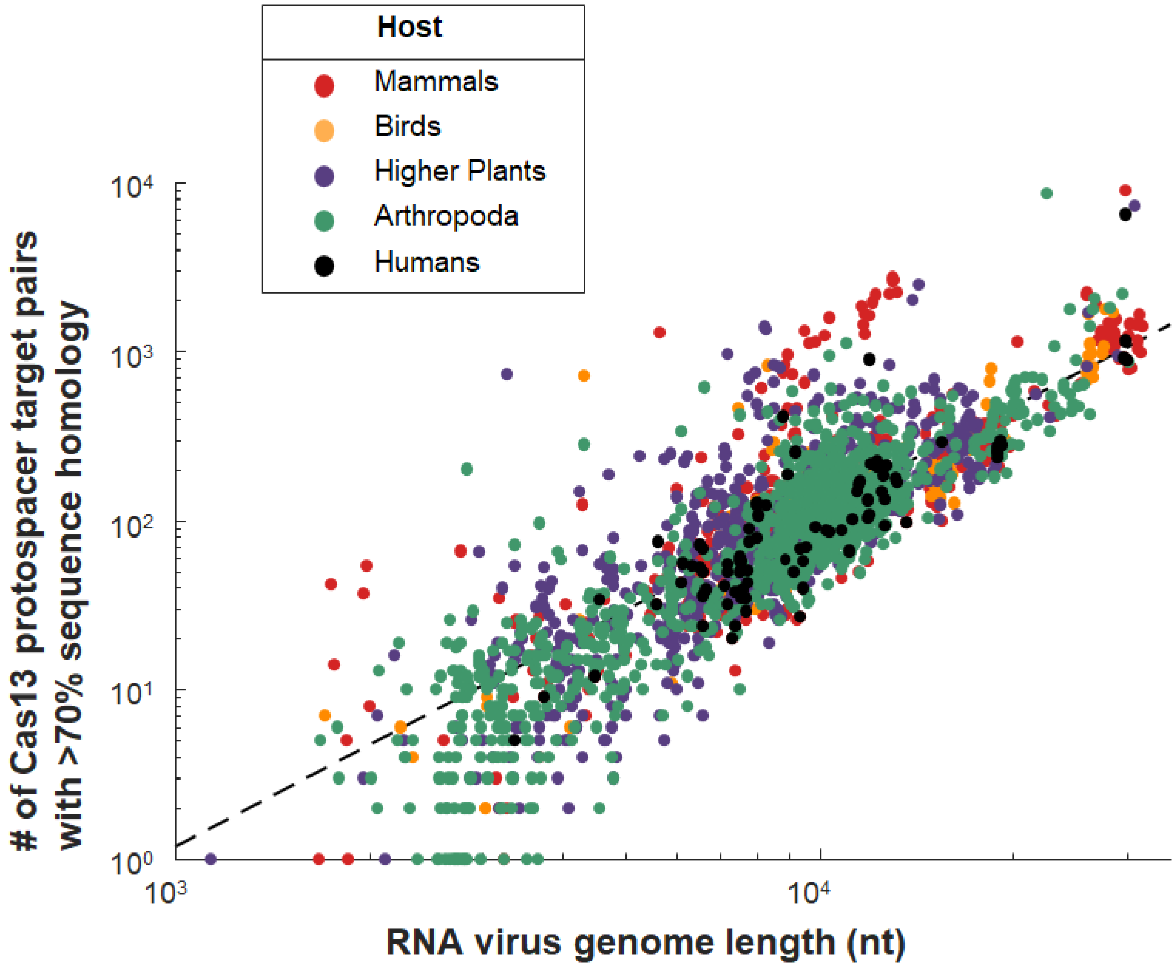


**Figure S2.** Pairs of targetable sites for Cas13 (23 nt), which share at least 70% homology, are abundant across the genomes of RNA viruses. The number of target pairs (P) vs. genome length (L) is fit well ($R^2$ = 0.7288) by: $\log_{10}(P) = 2.202 * \log_{10}(L) - 6.76$, or equivalently $P = 1.74e\text{-}7 * L^{2.202}$. All complete, RefSeq-quality genomes of RNA viruses, excluding proviruses, available by December 27, 2020 were downloaded from the NCBI Virus database with hosts: arthropoda (1074 viral species), aves (111), mammal (496), higher plant / embrophyta (691), and human (89). Genomes composed of multiple segments or CDS from the same viral isolate were considered together.

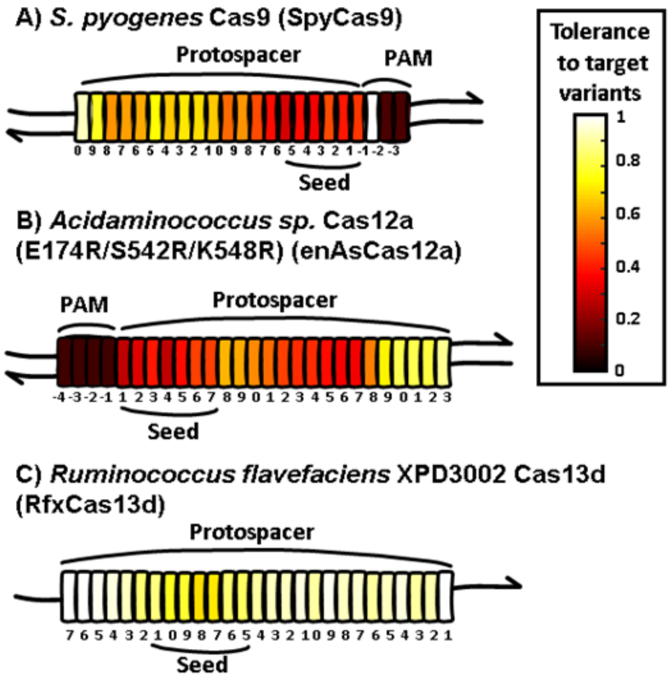


**Figure S3.** Position-dependent effects of sequence variants in the targeted region on the activity of different CRISPR effectors: (A) type II CRISPR effector Cas9 from *S. pyogenes* [5, 18], which targets dsDNA; (B) an engineered variant of type V CRISPR effector Cas12a from *Acidaminococcus sp*. BV3L6, enAsCas12a [3, 4], which targets dsDNA; and (C) type VI CRISPR effector Cas13d from *Ruminococcus flavefaciens* XPD3002 [2, 25], which targets ssRNA. Targeted nucleotides are coloured by the tolerance, or the average change in activity if there is a sequence variation at that site, of the CRISPR effectors for each position within or near the targeted (protospacer) sites. Tolerance of 1 implies that the CRISPR effector exhibits no change in activity regardless of the nucleotide identity at that site, and a tolerance of 0 implies that CRISPR activity is completely abolished if the nucleotide sequence is changed from the expected nucleotide at that site. The “seed” region is defined as a region of high sensitivity (low tolerance) to sequence variations at those sites; Cas9 and Cas12a are also highly sensitive to sequence variations at the protospacer adjacent motif (PAM) that are recognized by the enzyme itself rather than the gRNA. For the design of polyvalent gRNAs, we seek to maximize activity of a single gRNA at multiple viral sites by exploiting well-tolerated mismatch- and position- specific mispairings of the CRISPR effectors to minimize potential reductions of activity at these different sites.

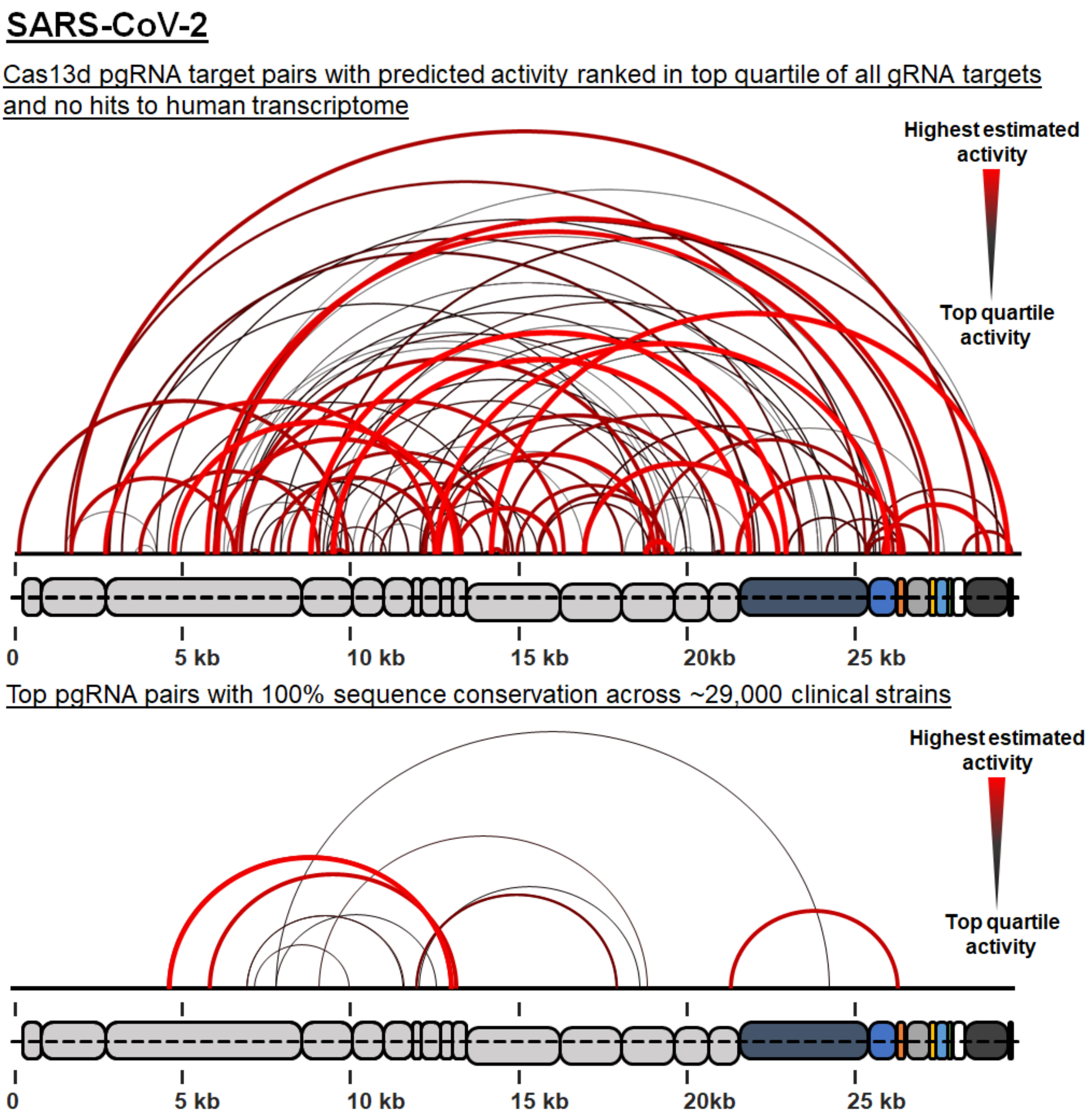


**Figure S4.** Target pairs in the SARS-CoV-2 genome for pgRNAs, 144 pairs of which predicted to have activity ranked in the top quartile of all anti-SARS-CoV-2 gRNAs at both sites and no predicted reactivity with the human transcriptome (top), 15 of which (bottom) target sites both with 100% sequence conservation across ~29,000 sequenced clinical variants.

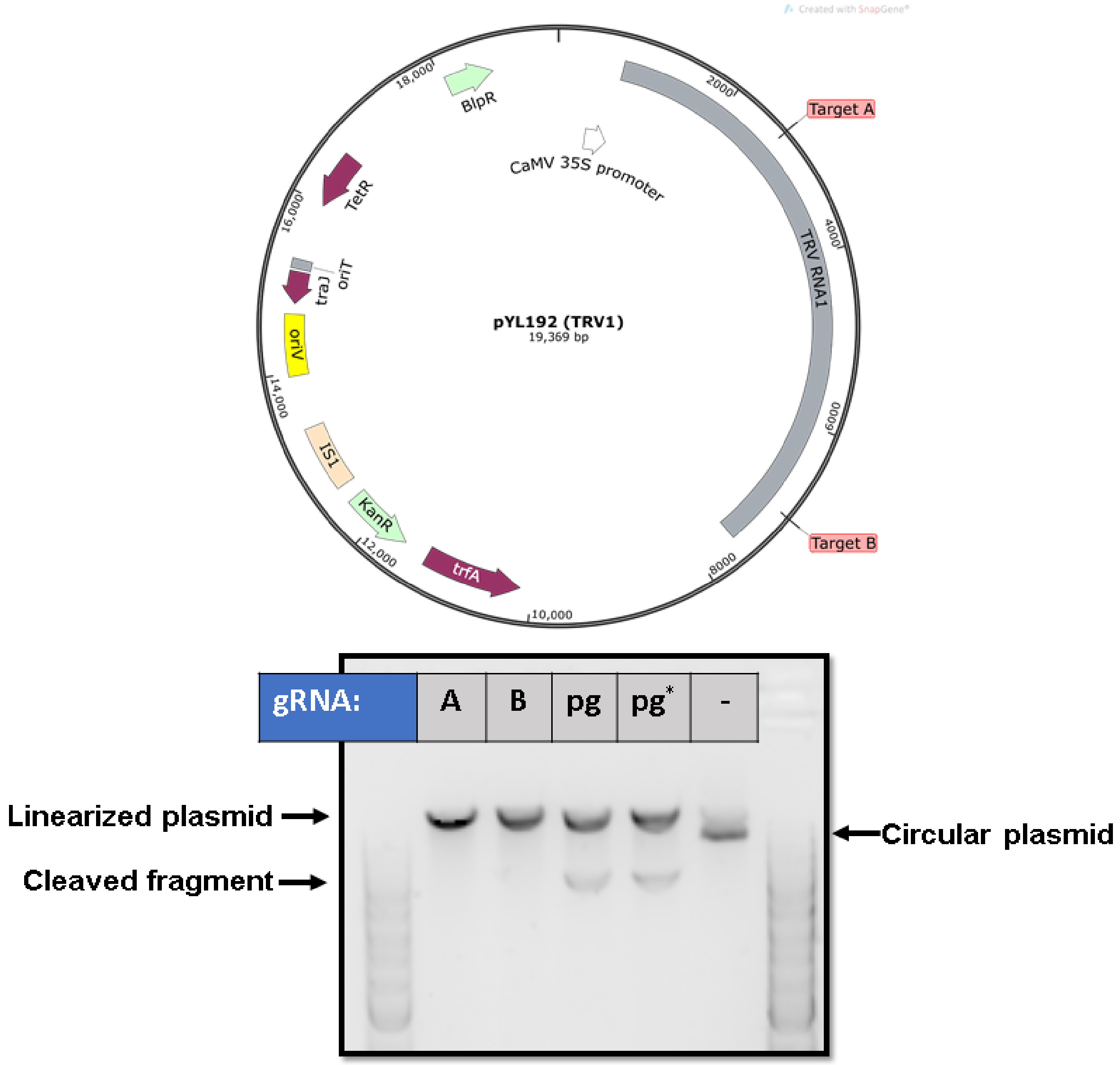


**Figure S5.** Simultaneous cleavage of two targets on the same plasmid by a Cas9-pgRNA RNP *in vitro*. (Above) Plasmid map with targets (see Figure 2) in red are located ~4kbp apart. (Below) Agarose gel electrophoresis after incubation of the plasmid and Cas9 with a "monovalent" gRNA specific to target A (A), a "monovalent" gRNA specific to target (B), or a "polyvalent" gRNA (pgRNA) optimized for activity at both targets. The sample marked with an * was incubated with and additional 1% polyethylene glycol during the cleavage reaction.

| Target | T. 7 | T. 7 | T. 8 | T. 8 | uncut | T. P. | T. 7 | T. 8 | T. 5 | T. 6 | T. 3 | T. 4 | T. 1 | T. 2 |
|---|---|---|---|---|---|---|---|---|---|---|---|---|---|---|
| gRNA | gRNA7 | gRNA8 | gRNA8 | gRNA7 | - | pgRNA | pgRNA | pgRNA | pgRNA | pgRNA | pgRNA | pgRNA | pgRNA | pgRNA |
| % Cleaved | 75.1% | 0% | 74.%5 | 0% | n/a | 78.2% | 0% | 88.2% | 32.4% | 79.7% | 70.5% | 72.7% | 71.3% | 63.5% |

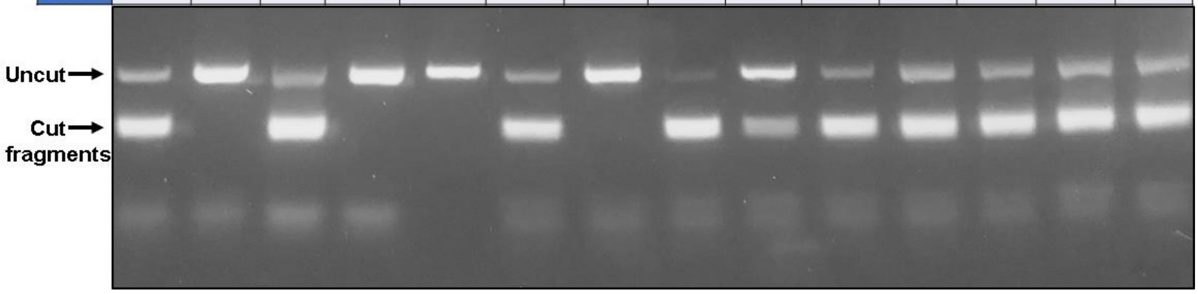


**Figure S6.** Agarose gel electrophoresis of Cas9-pgRNA cleavage products of synthetic targets with increasing divergence; related to Figure 2D. Sequences can be found in Supplementary Table S5.

| Target | HIV 4794 | HIV 9078 | HPV16 2716 | HPV16 7434 | HTLV1 4382 | HTLV1 7503 | HAvC 9382 | HAvC 11125 | HPV16 4525 | HPV18 934 | HAvC 731 | HAvC 25247 | HIV 4794 |
|---|---|---|---|---|---|---|---|---|---|---|---|---|---|
| gRNA | pgRNA-HIV. 4794s. 9078s | pgRNA-HIV. 4794s. 9078s | pgRNA-HPV16. 2716s. 7434a | pgRNA-HPV16. 2716s. 7434a | pgRNA-HTLV1. 4382a. 7503a | pgRNA-HTLV1. 4382a. 7503a | pgRNA-HAvC. 9382a. 11125a | pgRNA-HAvC. 9382a. 11125a | pgRNA-HPV16. 4525a. HPV18. 934s | pgRNA-HPV16. 4525a. HPV18. 934s | pgRNA-HAvC. 731s. 25247s | pgRNA-HAvC. 731s. 25247s | - |
| % Cleaved | 26.6% | 46.2% | 43.9% | 61.3% | 44.9% | 0% | 0% | 47.1% | 62.1% | 0% | 53.4% | 64.5% | n/a |

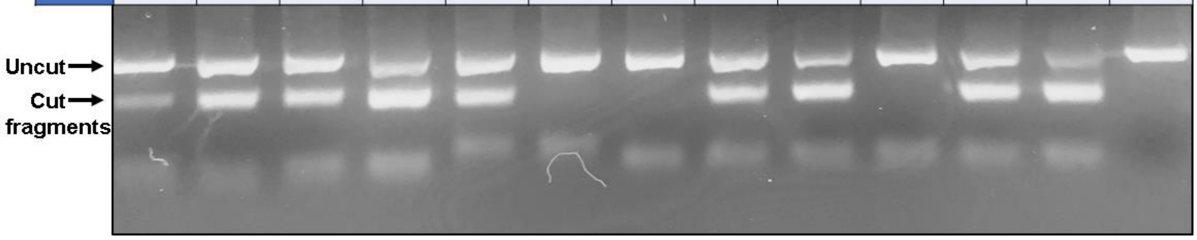


**Figure S7.** Agarose gel electrophoresis of Cas9-pgRNA cleavage products of synthetic targets with increasing divergence; related to Figure 2E. Sequences can be found in Supplementary Table S5.

| Target | A | A | A | B | B | B | A | A | A | B | B | B | A | A | A | B | B | B |
|---|---|---|---|---|---|---|---|---|---|---|---|---|---|---|---|---|---|---|
| gRNA | gA | gA | gA | gA | gA | gA | gB | gB | gB | gB | gB | gB | pg | pg | pg | pg | pg | pg |
| gRNA structure | sg | dupl. | dupl. | sg | dupl. | dupl. | sg | dupl. | dupl. | sg | dupl. | dupl. | sg | dupl. | dupl. | sg | dupl. | dupl. |
| 5' G | + | - | + | + | - | + | + | - | + | + | - | + | + | - | + | + | - | + |
| % Cleaved | 87 | 74 | 85 | 0 | 0 | 0 | 0 | 0 | 0 | 86 | 72 | 81 | 59 | 82 | 81 | 34 | 61 | 35 |

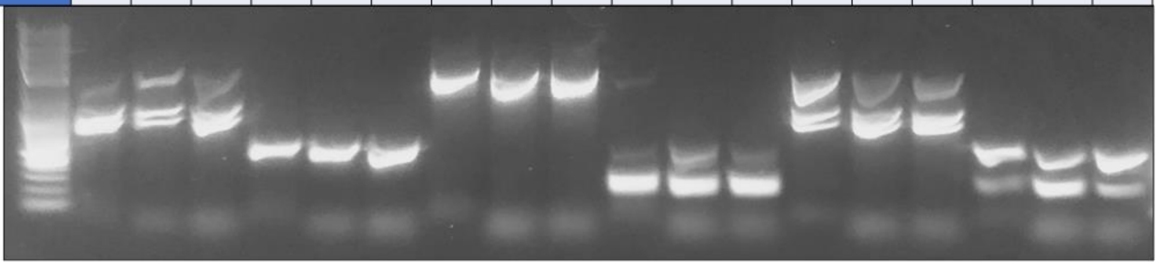

**Figure S8.** Agarose gel electrophoresis of Cas9-gRNA cleavage products of TRV targets (Figure 2A-C) under different *in vitro* conditions typically optimized for gene editing. Those gRNAs marked with 5'- have 21 nt long spacers with an unpaired 5'- G. Those with 'sg' gRNA structures were transcribed *in vitro* as 'single guides RNAs' (sgRNA) that fuse the crRNA and tracrRNA in continuous RNA molecule, while for 'dupl.' (duplex) the crRNA was synthetized (IDT; Coralville, IA) and hybridized with a tracrRNA (IDT) prior to incubation with the Cas9. Sequences can be found in Supplementary Table S5.

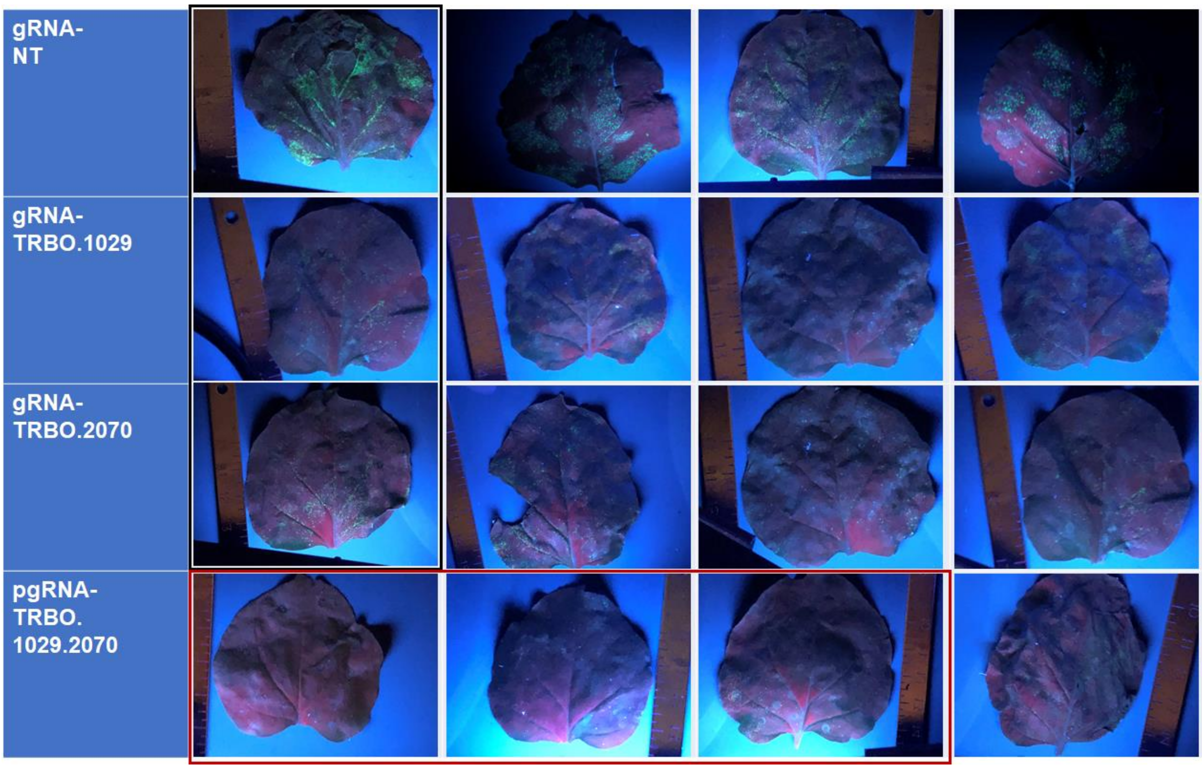


**Figure S9.** Additional photographs of leaves after induction of TRBO-GFP proliferation and transient expression of RfxCas13d with different gRNAs or pgRNAs. Related to Figures 3B and 3C, where images boxed in black and red are the same images as shown in Figure 3B in their respective colors.
